# Physiological and molecular mechanisms regulated mesophyll conductance under severe drought in water-saving drought-resistant rice

**DOI:** 10.1101/2022.03.24.485731

**Authors:** Haibing He, Lele Wang, Xuelan Zhang, Li Zhan, Quan Wang, Ru Yang, Cuicui You, Jian Ke, Liquan Wu

## Abstract

Water-saving and drought-resistant rice (WDR) is a new type of rice varieties. It plays an important role in responding to drought with high yield and has been widely planted in central China at present. High photosynthetic production potential caused by high mesophyll conductance (*g*_m_) is the main factor promoted high yield formation in drought for WDR. But little is known about physiological and molecular mechanisms regulated *g*_m_ in drought for WDR. Therefore, WDR cultivar HY73 and drought-sensitive cultivar HLY898 were used for comparative studies with three irrigation regimes before applying severe drought treatment at heading to create different differential individuals of photosynthetic potential and *g*_m_. The results showed that cultivar HY73 had lower up-regulation different expression genes (DEGs) than cultivar HLY898 in drought at transcriptional level. Conversely, DEGs of down-regulation was higher in cultivar HY73 than cultivar HLY898. In addition, 3071 DEGs were clustered in 3 modules named Midnightblue (734 DEGs), Blue (921 DEGs), and Turquoise (1416 DEGs) in severe drought merged three irrigation regimes and both cultivars, which the modules had significant correlational relationship with *g*_m_ based on weighted gene co-expression network analysis (*P*<0.05). Only DEGs in midnightblue module were enriched in photosynthesis process and positively regulated *g*_m_ (*P*<0.05). The main biological process were photosynthesis (GO:0015979), light harvesting in photosystem I (GO:0009768), reductive pentose-phosphate cycle (GO:0019253), protein-chromophore linkage (GO:0018298), photosynthetic electron transport in photosystem I (GO:0009773), and photosystem II repair (GO:0010206). These results indicate that *g*_m_ and energy distribution in PSI and PSII systems could synergistic effect photosynthetic production potential in severe drought for rice plants. In the modules, the 18 most highly connected hub genes were screened using co-expression networks method. RT-PCR analysis indicated that *CSP41B*, *PGLP1A*, *LHCA5*, and *GSTU6* genes had a similar variation trend with *g*_m_ among treatments for both cultivar. *LHCA5* and *CSP41B* genes were significantly up-regulated in HY73 compared with HLY898 in drought (*P*<0.05). And the both genes locates in thylakoid membrane in photosystems. Therefore, *LHCA5* and *CSP41B* genes could be key genes to synergistically manage *g*_m_ and energy distribution in photosystems. Our results provide some new physiological and molecular mechanisms regulated *g*_m_ in severe drought for WDR.

## Background

Drought is one of the most serious abiotic stresses affecting growth and development of rice plants. In major rice producing countries of Asia, rice plants are often exposed to drought environment with high frequency at different growth stages, which have a serious threat to world’s food security (Barker *et al*., 2000; Wang *et al*., 2019a). With global warming, the requirements for irrigation water will increase in paddy fields around the world (Belder *et al*., 2004; Ding *et al*., 2017). Take China for example, By 2080, irrigation demands may increase to 20-90 mm by 2080 in central China. The increment of irrigation amount would be greater than 100 mm in Hubei, Anhui, and Jiangsu provinces which are major rice producing areas in central China (Ding *et al*., 2017). We are currently faced with 3×10^10^ m^3^ irrigation gaps because of water shortage in agricultural production in China (Wang *et al*., 2018a). Rice is the most water-consuming crop, the shortage caused by irrigation increment is bound to aggravate the drought degree of rice plants. Therefore, water-saving, drought-resistant, and high-yield rice is an urgent issue to be solved in typical drought-prone areas around the world.

The limitation of photosynthesis is one of the main reasons for decreasing grain yield in drought (He *et al*., 2014; 2021; Wang *et al*., 2021). The homeostasis mechanism of reactive oxygen species (ROS) is disrupted and ROS contents would be increased in drought due to the reduction of oxygen molecular to superoxide on the acceptor side of PSI called Mehler reaction (Christine and Shigeru, 2011; Singh *et al*., 2018). The activity of Ribulose-1,5-bisphosphate carboxylase/oxygenase and the regeneration of ribulose-1,5-disphosphate (RuBP) are then blocked in the situation (Wang *et al*., 2019b). Photosynthetic organs would be damaged by excess luminous energy in drought (Kirchhoff, 2018). Even the damage is irreversibly repaired when PSI system is injured (Scheller and Haldrup, 2011). Photosynthetic productivity potential is also not completely recovered to pre-stress level after rehydrating if the PSI system is damaged (Scheller and Haldrup, 2005). Finally, grain yield is decreased because of inadequate source supply capacities caused by low photosynthetic potential in drought (Chu *et al*., 2014; He *et al*., 2014;Wang *et al*., 2021).

In recent years, a new type of rice varieties named water-saving drought-resistant rice (WDR) has been released from China. Generally, WDR has high yield and high quality characteristics accompanied with great drought-resistant ability (Luo *et al*., 2011). WDR has been widely planted in central China’s such as Anhui, Jiangxi, Hunan and Hubei (Hou *et al*., 2021). It has even become mainly-planted cultivars in the typical seasonal drought regions around Anhui Province. Compared with high-yield rice varieties, the high-yield and drought-resistant characteristics of WDR are closely related to the high photosynthetic potential of leaves in drought (He *et al*., 2021; Wang *et al*., 2021). In drought, stomatal conductance (*g*_s_) is significantly reduced to prevent water loss within cellular. Meanwhile, mesophyll conductance (*g*_m_) is also usually declined in drought for rice plants (Lauteri *et al*., 2014; Ouyang *et al*., 2017; Wang *et al*., 2019c). Generally, decreased photosynthetic production can be ascribed to inadequate CO_2_ concentration at carboxylation sites limited by low CO_2_ diffusion capacities from air to carboxylation sites in drought (Flexas *et al*., 2004). However, for WDR, *g*_m_ is slight reduction compared with drought-sensitive variety, even if the *g*_s_ is significantly reduced when responded to seasonal drought at heading, especially in optimized water management regime before heading (He *et al*., 2021). Maintaining high *g*_m_ is a pivotal reason to obtain high photosynthetic production potential for WDR in drought.

*g*_m_ is mainly affected by structural factors including leaf thickness, cell wall characteristics, mesophyll cell characteristics and chloroplast ultrastructure and physiological factors consisted of aquaporin and carbonic anhydrase (Evans *et al*., 2009). Increasing the number of mesophyll cells per unit area can significantly improve *g*_m_ of rice plants (He *et al*., 2017). *g*_m_ has positively correlation with the surface area of chloroplast exposed to intercellular air spaces per unit mesophyll cell area (Sc/S), and is negatively related with the cell wall thickness (Xiong *et al*., 2017). The decrease of Sc/S is considered to be the key factor leading to the decrease of *g*_m_ and photosynthetic potential of rice plants in drought (Xiong *et al*., 2017). However, Ellsworth *et al*. (2018) suggest that Sc/S has little effect on *g*_m_ of rice plants.

The main factors regulated *g*_m_ are related to cell wall characteristics. The results suggest that the effect of mesophyll cell structures on *g*_m_ could be related to varieties characteristics and stress intensities (Tomás *et al*., 2013).

At present, some of research results of aquaporin gene function are still controversial in regulating *g*_m_. In the early stage, barley *HvPIP2;1* gene was introduced into rice plants by transgenic method, *g*_m_ was increased in *PIP2;1* over-expressed plants (Hanba *et al*., 2004). But Ding *et al*. (2019) had proved that *OsPIP2;1* have no significant effect on *g*_m_ in rice when the gene was knocked out by gene editing. In addition, Huang *et al*. (2021) found that there was no significant difference in photosynthetic rate and *g*_m_ between PIP knockout lines and wild type of rice plants in field conditions. However, *g*_m_ and photosynthetic rate of PIP knockout lines were significantly lower than that of wild type in pot conditions. The reason for the difference in aquaporin genes function may be closely related to the growth environment (Kromdijk *et al*., 2020; Huang *et al*., 2021). These results also confirmed that aquaporin genes may not be the main factor regulating *g*_m_ in some specific environments. Probably, structural and physiological functions in regulating *g*_m_ require the induction of other genes.

Increased aquaporin activities could compensate for the negative effect of adverse leaf structure on *g*_m_ in drought for conventional high-yield rice varieties (Ouyang *et al*., 2017; He *et al*., 2021), but for WDR, the function of aquaporin did not improve and g_m_ did not also significantly decrease in drought (He *et al*., 2021). The results suggest that WDR might be different adaptive mechanisms in structural, physiological, and molecular levels. Therefore, WDR and drought-sensitive varieties were chosen as experimental materials with three water treatments in this study. Severe drought was applied at heading. RNA-seq, weighted gene co-expression network analysis, and protein interaction analysis technologies were synthetically adopted to reveal physiological and molecular mechanisms about maintaining high *g*_m_ in drought. The intentions of this study were (1) assessment the differences in transcription level of WDR in responding drought compared with drought-sensitive variety; (2) analysis the biological characteristics regulated *g*_m_ in drought; (3) screen key genes regulated *g*_m_ in drought. The research would provide important theoretical bases for screening and breading new WDR cultivars with high *g*_m_ traits in serve drought. Meanwhile, this study would be very useful to guide practices with high photosynthetic production potential by improving *g*_m_ for WDR in response to drought.

## Materials and methods

### Experimental design

Pot experiment was conducted inside of the movable awning at comprehensive experimental station of Anhui agricultural university, Lujiang, Anhui province, China (31°48′N, 117°23′E). Diameter and height of the pots were 30 cm and 40 cm, respectively. Each pot was filled with 20 kg fine-grained paddy soil with 2.00 g total N per kilogram soil, 32.40mg organic matter per kilogram soil, 24.80 mg Olsen-P per kilogram soil, and 19.40 g total K per kilogram soil. The soil *pH* was 5.9. All pot were planted one plant with 21-day-old seedlings from the nursery trays at same day. 12 plants including one main stem and 11 tillers were kept per pot by artificially pruning tillering buds to avoid potential effect of different population (tiller numbers) on individual growth vigour. At drought sensitive period which remaining leaf primordium number range from 0.2 to 0.5, severe drought degree with −50 KPa soil water potential monitored at 20 cm soil layer were imposed for 7 days. 2-3 cm water layer were irrigated for all pots after ending the drought till harvest.

Cultivar Hanyou 73 (HY73) was chosen as water-saving and drought-resistant rice. The cultivar has a large cultivated area in central China, which is a type high yield, high quality, and high drought-resistant variety. Cultivar Huiliangyou898 (HLY898) was identified as high yield, great quality, and drought sensitive variety based on the results of our previous varieties screening. The both cultivars are *Indica* hybrid rice variety grown in China as single-season rice. Different irrigation regimes were adopt to create diversity individual plants in this study, three water treatments were set before severe drought. Traditional flooding irrigation with 2-3 cm water layer was considered as control treatments, i.e. the soil water potential was always maintained at 0 KPa marked with W1 treatment.Mild wetting-drying alternate irrigation was marked with W2 treatment, which supplementary irrigation was repeatedly applied when soil water potential was reached at −10 Kpa at 15 cm soil layer before severe drought. Severe wetting-drying alternate irrigation was marked with W3 treatment, which supplementary irrigation was repeatedly applied when soil water potential was reached at −30 Kpa at 15 cm soil layer before severe drought.

All parameters were observed at pre-severe drought and the 7^th^ day of drought for both cultivars and three irrigation regimes. Each treatment had three biological repetition. Some monitored parameters such as gas exchange parameters are sensitivity to air environment. Therefore, the pots were moved to the artificial climate chamber to control some necessary environmental factors. Generally, daytime temperature, night temperature, relative humidity, photoperiod, and light intensity were set as 25 °C, 18 °C, 75%, 10 h, and 1200 μmol m^−2^ s^−1^, respectively.

## Measurements

### Gas exchange parameters

Net photosynthetic rate (*P*n) and stomatal conductance (*g*_s_) of flag leaves at steady state were determined by open-low gas exchange system Li-Cor 6400XT (Li-Cor Inc.) under 1200 μmol m^−2^ s^−1^ light intensity. CO_2_ concentration within chamber was 400 μmol mol^−1^ supplied by a CO_2_ cylinder. Leaf-to-air vapor pressure deficit was averagely controlled at 1.3 KPa.

### Calculation of *g*_m_

Farquhar-von Caemmerer-Berry model was used to determine mesophyll conductance (*g*_m_). The chosen model was modified by Harley *et al*. (1992). It can be described as following.

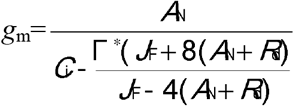

Γ* is the compensation point of CO_2_ without the day respiration. R_d_ is respiration rate during the day. *J*_F_ represents electron transport rate in PSII system. The parameters were obtained by referring to our previous study (He *et al*., 2021).

### RNA-seq analysis

The samples were extracted for total RNA and then subjected to RNA quality control. The quality standards of the library are as follows: agarose gel electrophoresis was used to detect the presence of DNA contamination and RNA integrity. The purity and integrity of RNA were accurately detected at OD260/280 and OD260/230 ratios. The constructed library was sequenced by Illumina HiSeqTM.

The raw reads obtained from sequencing were subjected to a series of cuts and rejects to obtain High-quality clean reads, which were homogenized and converted into FPKM values (Fragments per kilobase of transcript per Million mapped reads). The differentially expressed genes (DEGs) were screened according to FDR<0.05 and |log2FC|>2, and three sets of differentially expressed genes were obtained for subsequent analysis of GO, KEGG, and transcription factor annotation.

### WGNA analysis

Weighted gene co-expression network analysis (WGCNA) is an algorithm for mining module (module) information from high-throughput expression data. In this method, a module is defined as a set of genes with similar expression profiles. If some genes always have similar expression changes in a physiological process or different tissues, then it is reasonable to assume that these genes are functionally related and can be defined as a module.

WGCNA analysis includes the following analysis processes: (1) Raw data processing. The raw data totaled 37358 genes with 18 samples, and genes with low fluctuations in expression variation (standard deviation≤0.45) were filtered. (2) Network construction parameters, setting power values from 1-30 and calculating the correlation coefficients corresponding to the networks they obtained and the average connectivity of the networks, respectively. (3) Module identification and analysis, based on the selected power values, a weighted co-expression network model was built. (4) Module trait association analysis, using the Pearson correlation algorithm to calculate the correlation coefficients and p-Value of module trait genes with physiological phenotypic traits such as *P*_n_, *g*_s_ and *g*_m_.(5) Core genes analysis. The top 50 genes with the highest connectivity in each module were analyzed to show the relationship between these genes, which the method were usually used to mine the core genes.

### Protein interaction analysis

For the core genes obtained based on transcriptome and WGCNA analysis methods, protein interactions were established for water channel protein genes, carbonic anhydrase genes and cell wall metabolism-related genes, which are closely related to the regulation of gm conductivity. The analysis was performed using String online database and Cytoscape was used to map the interactions.

### qRT-PCR verification of candidate genes

To check the accuracy of sequencing results, 15 candidate genes were sequenced in this study, and the leaf samples of each species before and after drought treatment were used for qRT-PCR quantitative analysis. (1) RNA extraction: total RNA was extracted from leaves and seeds using the RNA Extraction Kit (DP432), and its purity was detected at 260 nm with a spectrophotometer, and samples with an A260/A280 reading of 1.8-2.1 could be used for further analysis, while the integrity of RNA samples could be detected by ordinary agarose gel electrophoresis. (2) cDNA preparation: reverse transcription using the cDNA synthesis kit and following the instructions(Takara, PrimeScript^Tm^ RT Master Mix). (3) determination of the transcript levels of the target genes with the Applied Biosystems 7300 fast real-time PCR System. The sequence numbers and sequences of the differentially expressed gene primers are shown in Table 1S.

### Statistical analyses

Differences between means for the *P*_n_, *g*_s_, and *g*_m_ parameters were compared by Fisher’s least significant difference (LSD) test at 5% significant level in SPSS16.0. Linear regression model and Pearson correlation analysis were used to reveal inherent relationships among *g*_m_ and candidate genes using in SPSS 16.0. Figures were constructed using OriginPro 8.5 software for the parameters. R package was used for creating KEGG, GO, genes clustering in modules, and protein interaction network diagrams.

## Results

### *P*_n_, *g*_s_, and *g*_m_ of WDR under severe drought

Cultivar HY73 had an apparently higher *P*_n_, *g*_s_, and *g*_m_ than cultivar HLY898 in the all water treatments (Fig.1). The observed parameters were decreased in severe drought than in sufficient irrigation for the three water treatments. Generally, there were greater decreasing amplitude in cultivar HLY 898 (decline by 30.71~50.02% and 30.77~41.67% for *P*_n_ and *g*_m_, respectively) than that in cultivar HY73 (decrease by 22.22~39.44% and 5.26~35.71% for *P*_n_ and *g*_m_, respectively) for *P*_n_ and *g*_m_ in severe drought compared with sufficient irrigation (Fig. 1A and C). However, *g*_s_ was reduce by 57.69~ 68.89% in cultivar HY73 in severe drought when compared with sufficient irrigation, which the falling range was 1.37~6.89 times higher for cultivar HY73 than for cultivar HLY898 (Fig. 1B). For water treatments, the W2 treatment had the greatest values for all of measured parameters, and then followed by the W1 and W3 treatments across both cultivars in both observed periods (Fig. 1).

**Fig. 1.**
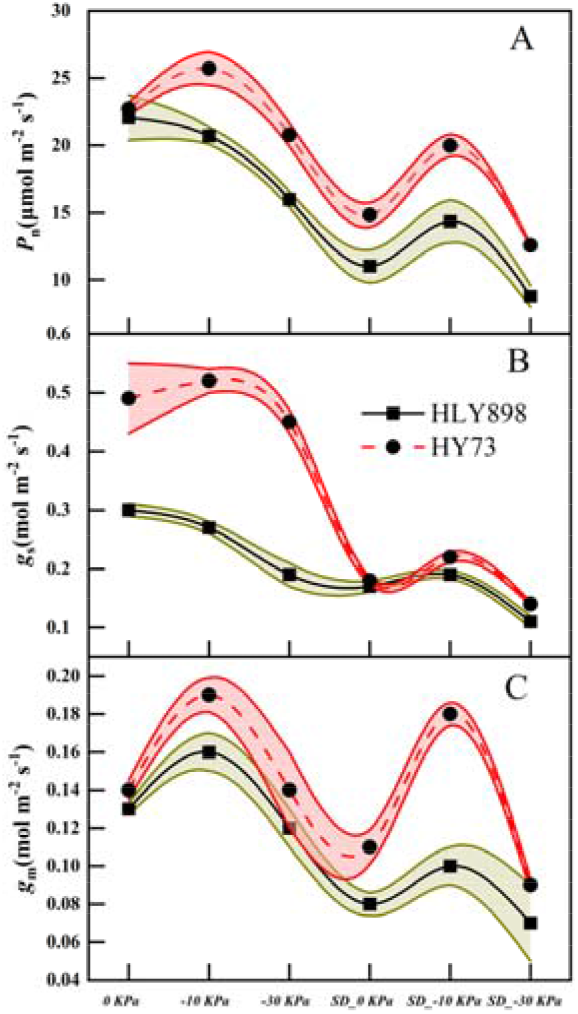
*P*_n_ (A), *g*_s_ (B), and *g*_m_ (C) of cultivars HLY898 (black-solid line) and HY73 (red-dotted line) in different water treatments. 0 KPa, −10 KPa, and-30 KPa represent the treatments of sufficient supplementary irrigation. Also, SD_0 KPa, SD_-10 KPa, and SD_-30 KPa represent the three water treatments of 0 KPa, −10 KPa, and −30 KPa in severe drought, respectively. The filled areas are standard deviations of different water treatments in cultivar HLY898 (filled with red part) and cultivar HY73 (filled with red dark yellow part). Mean values and standard deviation were obtained from three replicates (n=3).

### Analysis of differentially expressed genes

The referenced transcriptome sequencing of 36 samples were completed in sufficient irrigation and severe drought. After removing low-quality reads, rRNA reads, reads containing adapters, and reads with > 10 unknown nucleotides, the clean data ranged from 6.22 ×10 ^9^ to 6.55 ×10 ^9^ in sufficient irrigation and from 6.53×10 ^9^ to 6.73 ×10 ^9^ in severe drought across water treatments and cultivars (Table 1). After filtering, Q30 was 90.44~91.40 % in sufficient irrigation treatments and 93.51~93.80 % in severe drought in three water treatment and two cultivars. The sufficient irrigation had 44.90×10 ^6^~48.49×10 ^6^ clean reads, and severe drought treatment had 47.91×10 ^6^~49.01×10 ^6^ clean reads for three water treatments of two cultivars (Table 1).

**Table 1.**
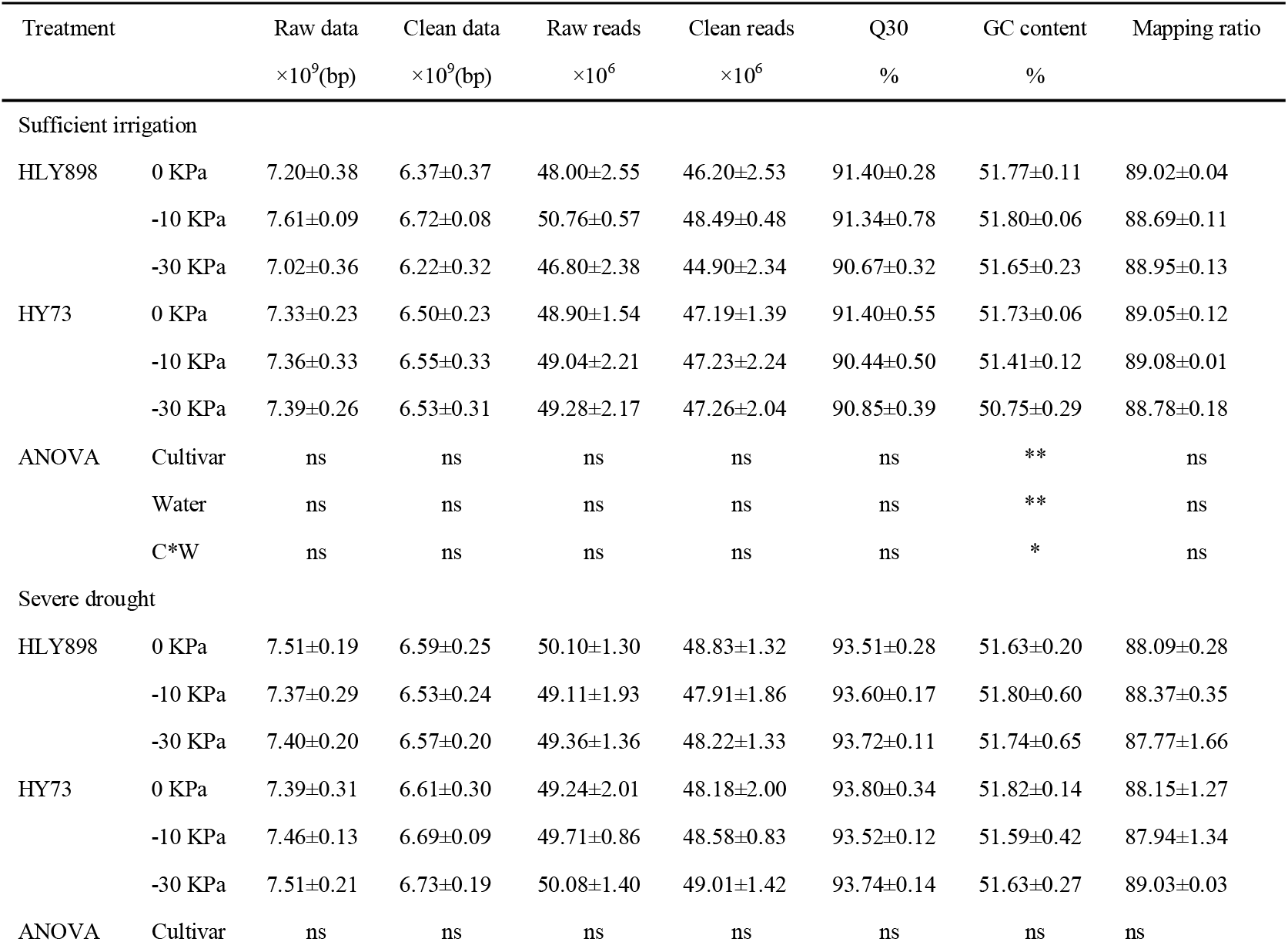
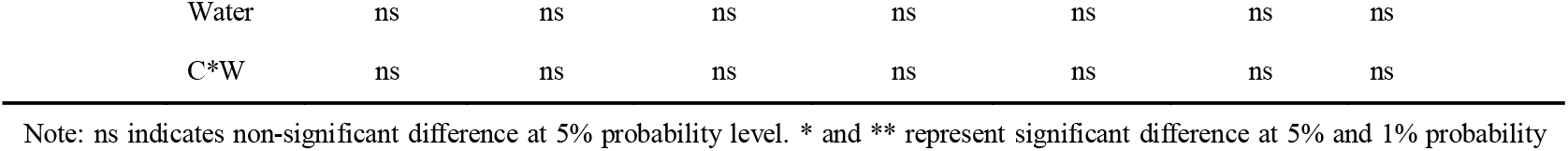
Summary of the transcriptome sequencing data and qualify filtering

| Treatment |  | Raw data | Clean data | Raw reads | Clean reads | Q30 | GC content | Mapping ratio |
| --- | --- | --- | --- | --- | --- | --- | --- | --- |
| | | $\times 10^9$(bp) | $\times 10^9$(bp) | $\times 10^6$ | $\times 10^6$ | % | % | |
| Sufficient irrigation |  |  |  |  |  |  |  |  |
| HLY898 | 0 KPa | 7.20±0.38 | 6.37±0.37 | 48.00±2.55 | 46.20±2.53 | 91.40±0.28 | 51.77±0.11 | 89.02±0.04 |
|  | -10 KPa | 7.61±0.09 | 6.72±0.08 | 50.76±0.57 | 48.49±0.48 | 91.34±0.78 | 51.80±0.06 | 88.69±0.11 |
|  | -30 KPa | 7.02±0.36 | 6.22±0.32 | 46.80±2.38 | 44.90±2.34 | 90.67±0.32 | 51.65±0.23 | 88.95±0.13 |
| HY73 | 0 KPa | 7.33±0.23 | 6.50±0.23 | 48.90±1.54 | 47.19±1.39 | 91.40±0.55 | 51.73±0.06 | 89.05±0.12 |
|  | -10 KPa | 7.36±0.33 | 6.55±0.33 | 49.04±2.21 | 47.23±2.24 | 90.44±0.50 | 51.41±0.12 | 89.08±0.01 |
|  | -30 KPa | 7.39±0.26 | 6.53±0.31 | 49.28±2.17 | 47.26±2.04 | 90.85±0.39 | 50.75±0.29 | 88.78±0.18 |
| ANOVA | Cultivar | ns | ns | ns | ns | ns | ** | ns |
|  | Water | ns | ns | ns | ns | ns | ** | ns |
|  | C*W | ns | ns | ns | ns | ns | * | ns |
| Severe drought |  |  |  |  |  |  |  |  |
| HLY898 | 0 KPa | 7.51±0.19 | 6.59±0.25 | 50.10±1.30 | 48.83±1.32 | 93.51±0.28 | 51.63±0.20 | 88.09±0.28 |
|  | -10 KPa | 7.37±0.29 | 6.53±0.24 | 49.11±1.93 | 47.91±1.86 | 93.60±0.17 | 51.80±0.60 | 88.37±0.35 |
|  | -30 KPa | 7.40±0.20 | 6.57±0.20 | 49.36±1.36 | 48.22±1.33 | 93.72±0.11 | 51.74±0.65 | 87.77±1.66 |
| HY73 | 0 KPa | 7.39±0.31 | 6.61±0.30 | 49.24±2.01 | 48.18±2.00 | 93.80±0.34 | 51.82±0.14 | 88.15±1.27 |
|  | -10 KPa | 7.46±0.13 | 6.69±0.09 | 49.71±0.86 | 48.58±0.83 | 93.52±0.12 | 51.59±0.42 | 87.94±1.34 |
|  | -30 KPa | 7.51±0.21 | 6.73±0.19 | 50.08±1.40 | 49.01±1.42 | 93.74±0.14 | 51.63±0.27 | 89.03±0.03 |
| ANOVA | Cultivar | ns | ns | ns | ns | ns | ns | ns |

|  |  |  |  |  |  |  |  |
| --- | --- | --- | --- | --- | --- | --- | --- |
| Water | ns | ns | ns | ns | ns | ns | ns |
| C*W | ns | ns | ns | ns | ns | ns | ns |

Note: ns indicates non-significant difference at 5% probability level. \* and \*\* represent significant difference at 5% and 1% probability level, respectively.

Meanwhile, GC content was 51.52% and 51.70% in sufficient irrigation and severe drought, respectively (Table 1). Among the 36 samples, the matching rates of clean read to the gene group were all greater than 87.77%. These results indicate that the sequencing data were of high quality.

In sufficient irrigation, up-regulated differential genes and down-regulated differential genes had a slight increasing trend with decreasing soil water potential when compared with traditional flooding irrigation treatment (0 KPa) across both cultivars. The number of differential genes were only 302~570 among different water treatments of both cultivar in sufficient irrigation. However, up-regulated differential genes and down-regulated differential genes were reached to 3178~3753 and 2365~3509 in severe drought when compared with 0 KPa treatment across three water treatments and cultivars (Fig. 2). Generally, there were more up-regulated genes in cultivar HLY898 than in cultivar HY73 after severe drought. But the down-regulated genes were more higher in cultivar HY73 than in cultivar HLY898 in severe drought compared to traditional flooding irrigation treatment. For cultivar HY73, up-regulated and down-regulation genes were increase with decreasing soil water potential in severe drought. But an opposite trend was observed for up-regulated genes in cultivar HLY898 among different water treatments, which decreasing soil water potential declined the number of differential genes for cultivar HLY898 (Fig. 2). The results shown that there could exist different response mechanisms to drought in both cultivars.

**Fig. 2.**
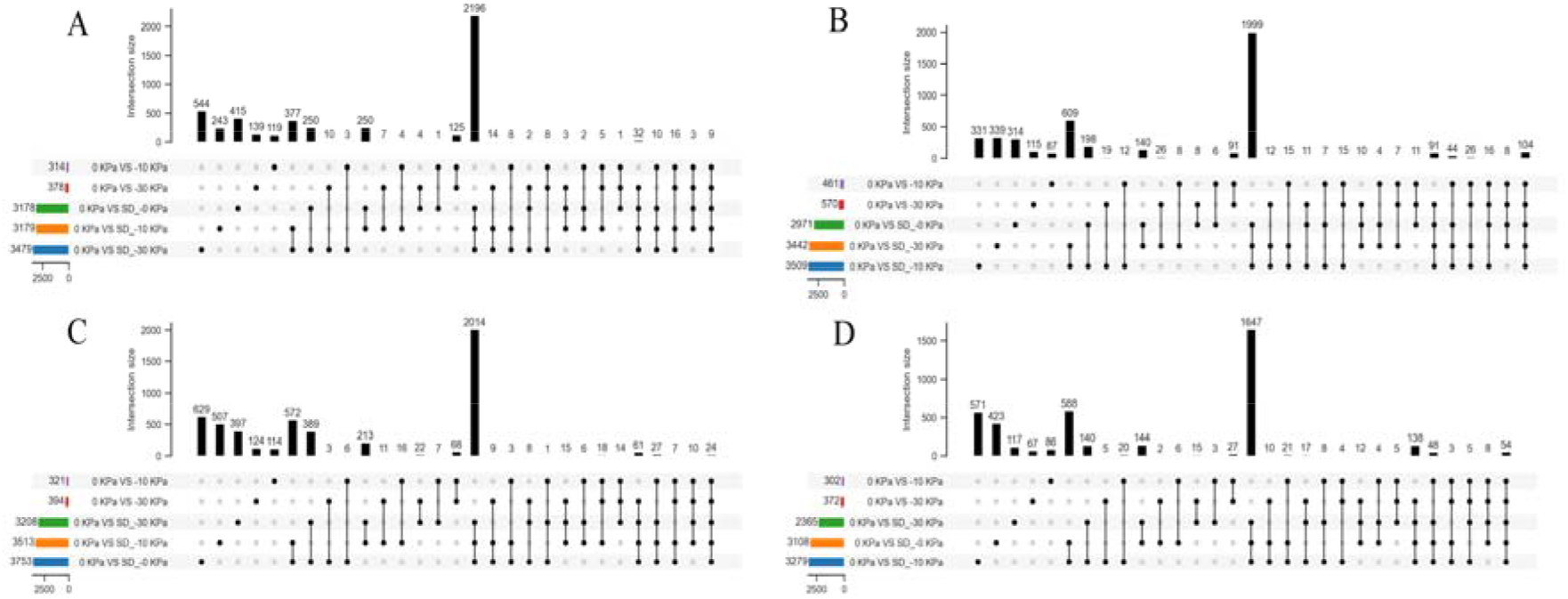
Up-regulated differential genes (A and C) and down-regulated differential genes (B and D) between different water treatments of cultivar HY73 (A and B) and cultivar HLY898 (C and D) at both observed periods. 0 KPa VS −10 KPa and 0 KPa VS −30 KPa indicates differential genes in the condition of - 10 KPa and −30 KPa when compared with 0KPa treatment in sufficient irrigation, respectively. Meanwhile, 0 KPa VS SD_0 KPa, 0 KPa VS SD_-10 KPa, and 0 KPa VS SD_-30 KPa represent differentially expressed genes of 0KPa, −10KPa, and −30 KPa treatments in severe drought compared with 0 KPa treatment in sufficient irrigation. The horizontal colored columns represent the total number of differentially expressed genes in each group. The vertical and black column represents the number of differentially expressed genes in each group corresponding to the black dots. The black line among/between the black dots indicates the same differentially expressed genes among/between different compared groups.

### Weighted gene correlation network analysis of differentially expressed genes

Weighted gene co-expression network analysis was severally adopted for the two observed periods of sufficient irrigation and severe drought. Each period had 18 samples assigned to two cultivars and three water treatments. 2778 genes and 5155 genes were selected for WGCNA analysis after filtering low variation (standard deviation≤0.5) in the expression matrix in sufficient irrigation and severe drought, respectively. Soft threshold was considered as 30 and 14 in sufficient irrigation and severe drought, respectively (Fig. 1S).

Three modules named brown, green, and red were screened in sufficient irrigation. There were 1322, 1048, and 81 genes in brown, green, and red modules, respectively (Fig. 3A). Brown module had significant negative correlation with *g*_s_ (r=−0.94^***^) and *P*_n_ (r=−0.60^**^). But green module had significant positive correlation with *g*_s_ (r=0.92^***^) and *P*_n_ (r=0.54^**^). Moreover, red module had significant positive correlation with *g*_s_ and *g*_m_ at 1% and 5% probability levels, respectively (Fig. 3A).

**Fig. 3.**
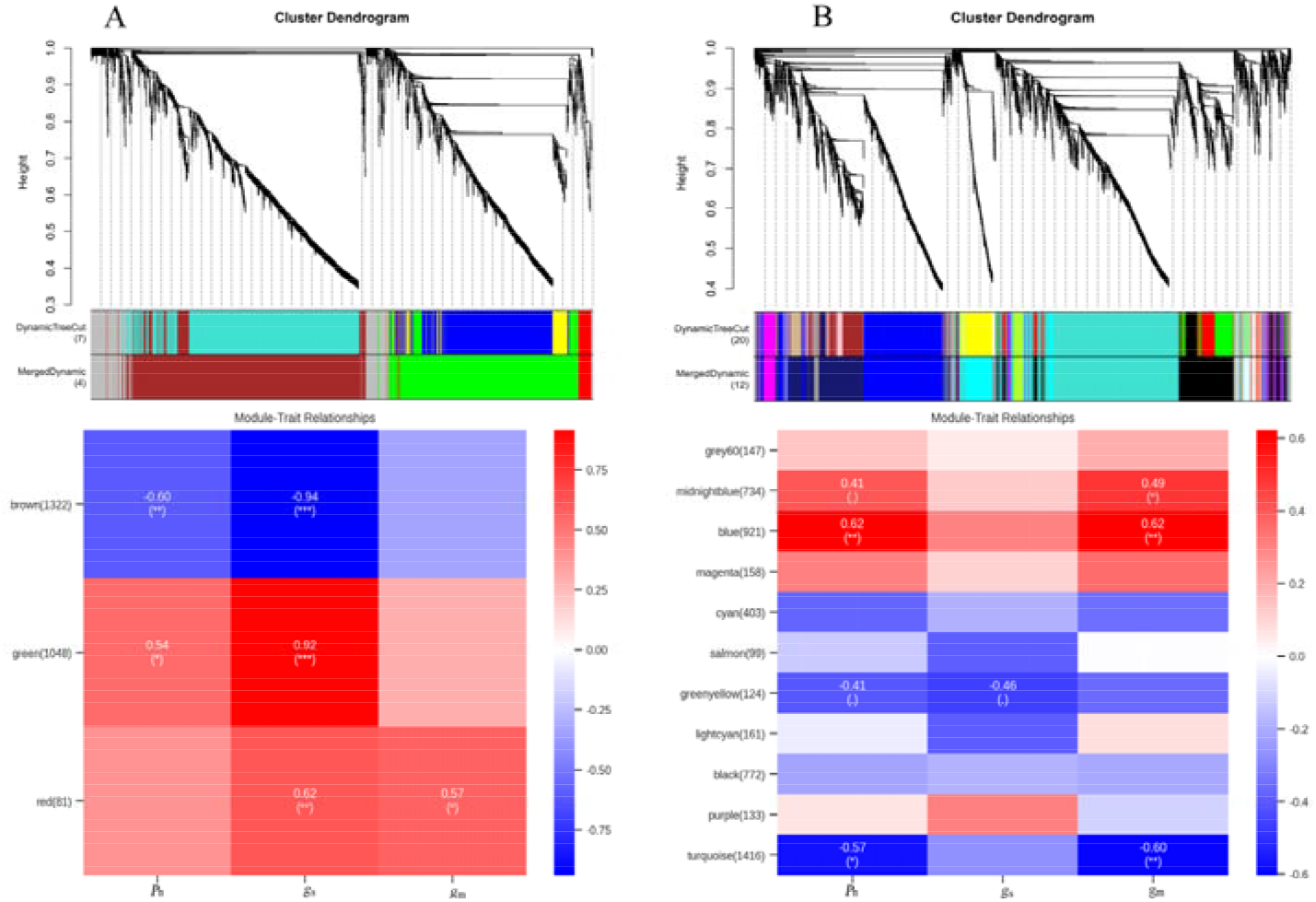
Gene co-expression network gene clustering number and modular cutting and its correlation analysis with *P*_n_, *g*_s_, and *g*_m_ in sufficient irrigation (A) and severe drought (B). *, **, and *** represent significant correlation between model and parameter at 5%, 1%, and 0.1% probability level.. indicates no significant correlational relationship between model and parameter.

There were 11 modules in severe drought. Turquoise module gathered the largest number of genes (1416) among all modules, which had significant negative correlation with *g*_m_ (r=−0.60^**^) and *P*_n_ (r=−0.57^*^). The second largest module contained 921 genes was blue. Blue module had significant positive correlation with *g*_m_ (r=0.62^**^) and *P*_n_ (r=0.62^**^). 734 genes was enriched in midnightblue module. The module was unique significant positive correlation with *g*_m_ at 5% probability level (r=0.49). The rest modules had no significant correlation with *P*_n_, *g*_s_, and *g*_m_ (Fig. 3B).

### Functional annotation of target modules

Although brown and green modules were significantly related with *g*_s_ and *P*_n_ in sufficient irrigation, but the GO and KEGG items were fewly enriched in photosynthesis or photosynthesis related pathways (Fig. 2S; Fig. 3S). The similar phenomenon was found in red module in sufficient irrigation (Fig. 4S). The results indicated that the enriched differentially expressed genes could not be directly regulation photosynthetic performance in sufficient irrigation.

In severe drought, biological process of top three items in blue modules were defense response to bacterium (GO:0042742; *P*=0.002), anthocyanidin reductase activity (GO:0033729; *P*=0.002), and regulation of jasmonic acid mediated signaling pathway(GO:2000022; *P*=0.005). But false discovery rate (FDR) values of the items were more than 0.05 (Fig. 4A). Correspondingly, KEGG pathways were mainly enriched in plant hormone signal transduction (Osa04075), plant-pathogen interaction (Osa04626), and sulfur metabolism (Osa00920) in blue modules (*P*<0.05; FDR>0.05; Fig. 4B). For the turquoise module, the main biological processes contained heme binding(GO:0020037), L-lysine transmembrane transporter activity(GO:0015189), and L-alanine transmembrane transporter activity (GO:0015180) in severe drought (*P*<0.05; FDR<0.05; Fig. 5A). Similarly, no related photosynthesis biological process and pathway were significantly annotated in turquoise module (Fig. 5). The results meant that blue and turquoise modules could directly contribute other metabolic pathways instead of photosynthesis pathways. And then indirectly affecting changes in *g*_m_ and *P*_n_.

**Fig. 4.**
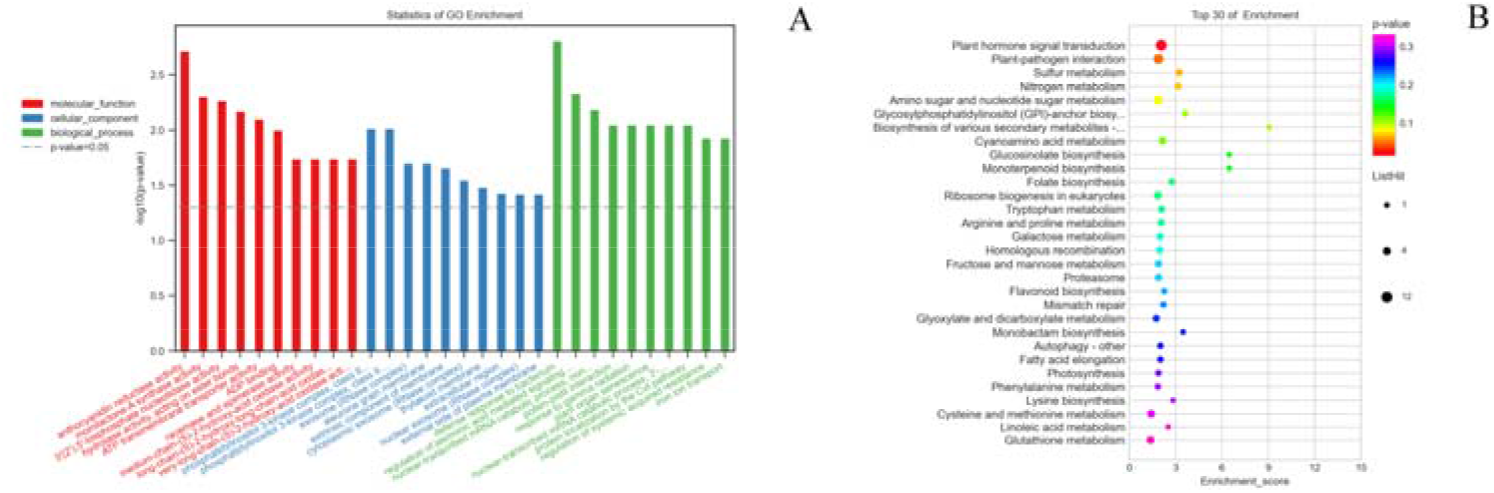
The first 10 GO (A) and 30 KEGG (B) items in the blue module based on WGCNA analysis in severe drought.

**Fig. 5.**
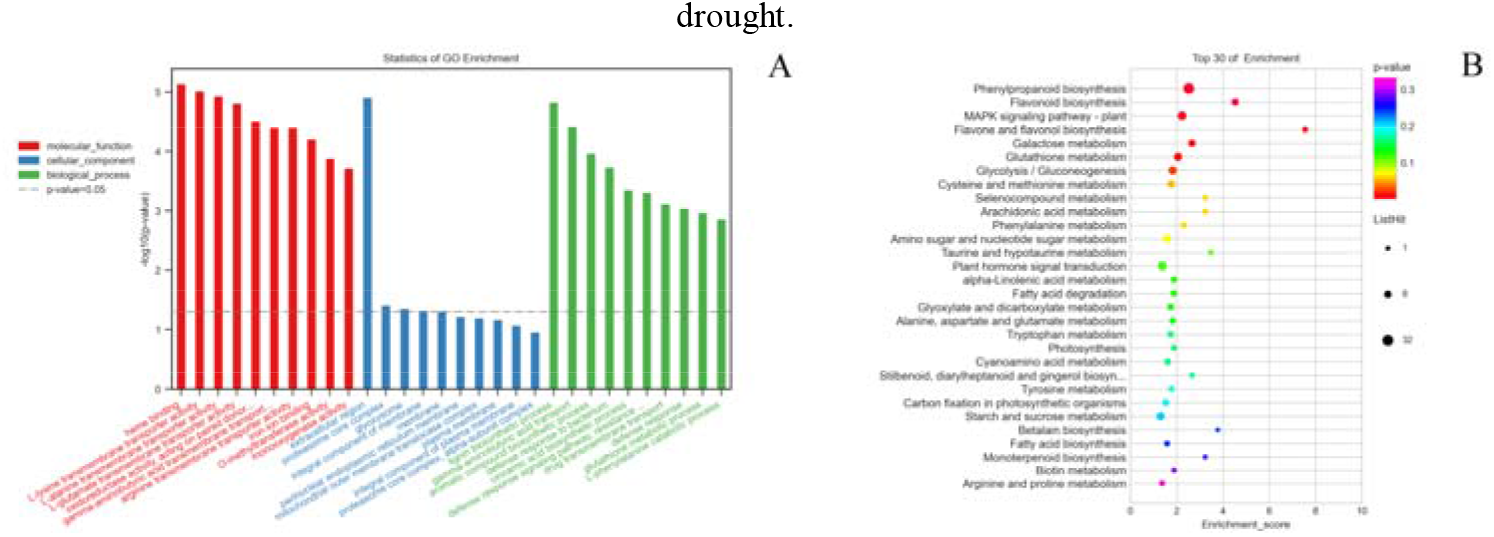
The first 10 GO (A) and 30 KEGG (B) items in the turquoise module based on WGCNA analysis in severe

In midnightblue module, both biological process and cellular component of enriched genes were significant relevant to photosynthesis (Fig. 6A; Table 2; *P*<0.05; FDR<0.05). For instance, biological process of top six were photosynthesis (GO:0015979), light harvesting in photosystem I (GO:0009768), reductive pentose-phosphate cycle (GO:0019253), protein-chromophore linkage (GO:0018298), photosynthetic electron transport in photosystem I (GO:0009773), and Photosystem II repair (GO:0010206) (Table 2). Meanwhile, the KEGG pathway was mainly and significantly enriched in photosynthesis process (Fig. 6B; *P*<0.05; FDR<0.05).

**Fig. 6.**
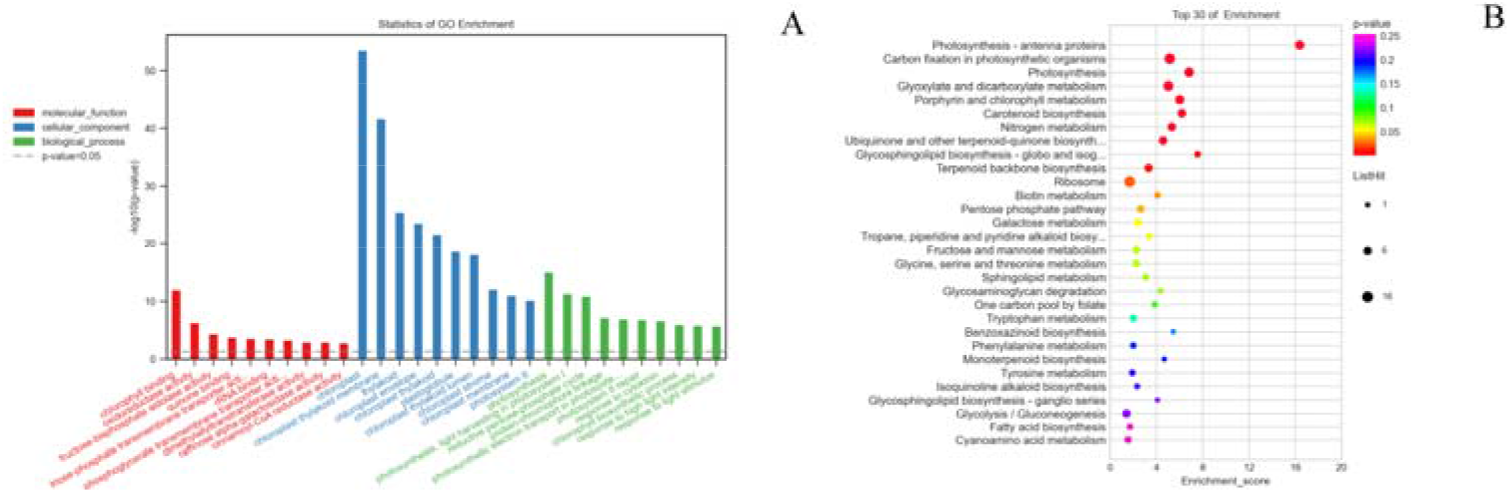
The first 10 GO (A) and 30 KEGG (B) items in the midnightblue module based on WGCNA analysis in severe drought.

**Table 2.**
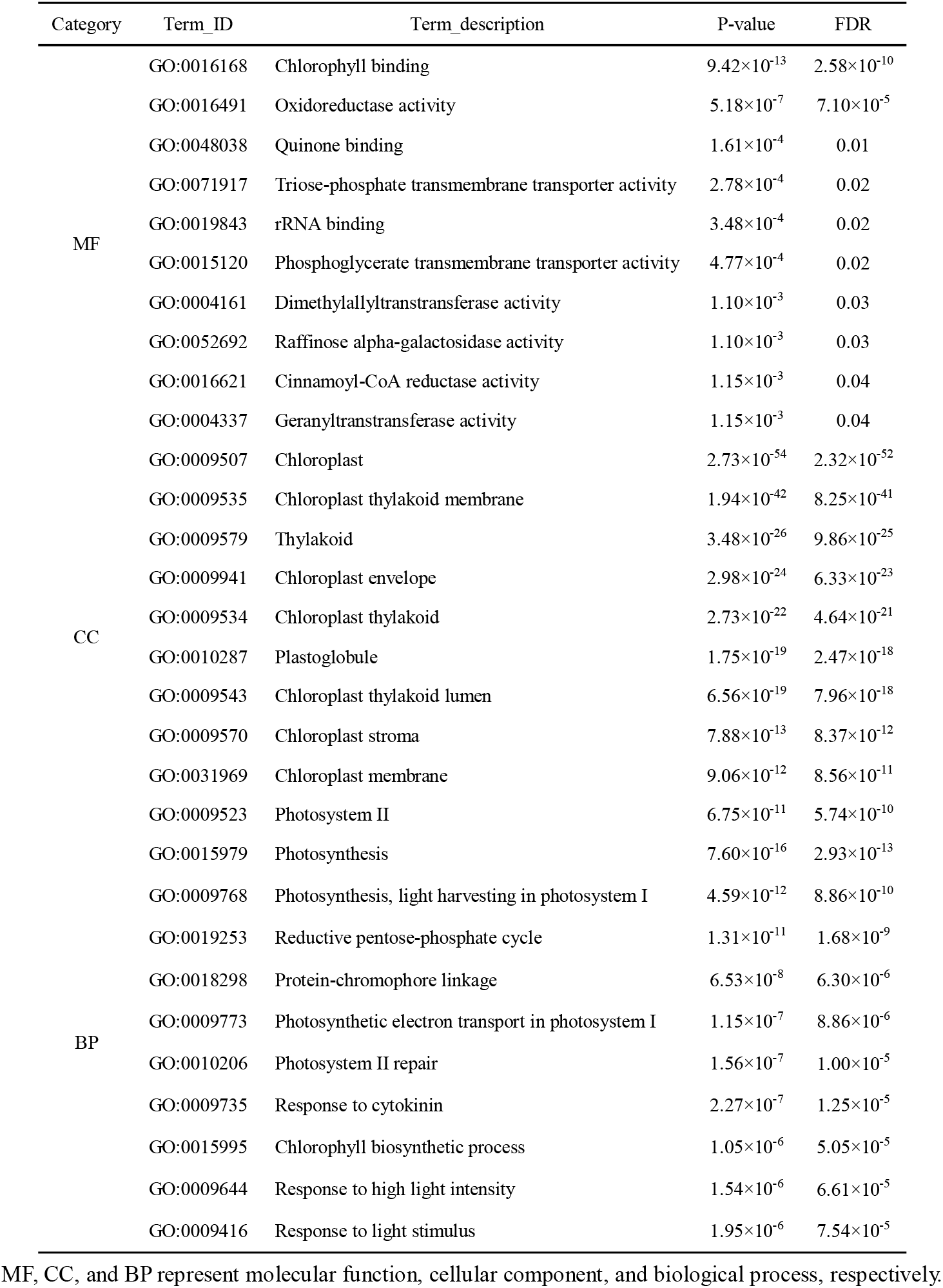
GO enrichments of midnightblue module (part)

### Hub genes identification of target modules

The core genes in a module were defined as the group of genes with the high connectivity between these genes in the module. There were 32 annotated genes in midnightblue (Fig. 7A). A lot of high connectivity genes associated with thylakoid functions, for instance *CSP41B*, *RBCS*, *PGLP1A*, *GAPA*, *DAAT*, *LHCA5*, *CAS*, *GSTU6*, *CHLP*, *TROL*, *At1g67280*, *CLEB3J9*, et al. were only found in the module (Fig. 7A). The connectivity of *SPQ*, *SSL*, and *SKL1* genes were the lowest among the genes.

**Fig. 7.**
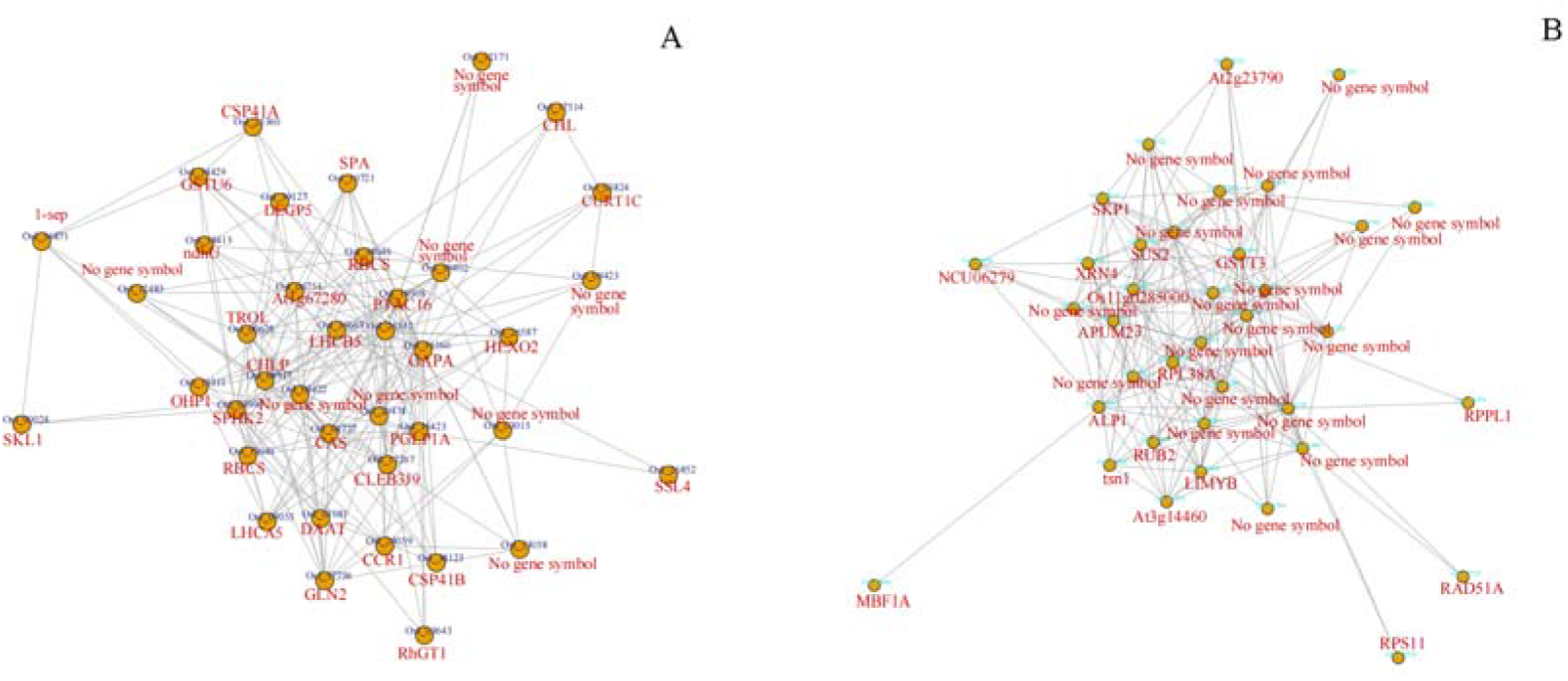
Co-expression networks of the most highly connected hub genes in the midnightblue module (A) and the turquoise module (B) in severe drought.

37 high connectivity genes were in turquoise module. Five co-expression genes such as *At2g23790*, *GSTT3*, *Apim23*, *LIMYB*, and *Os11g0285000* were found in turquoise and brown modules (Fig. 7B; Fig. 6S). There were 19 genes with no gene symbol in the genetic database in turquoise module (Fig. 7B).

36 high connectivity genes were obtained in blue module in severe drought (Fig. 5S–B). However, there were for 18 genes with no gene symbol in blue module. Meanwhile, 9 genes, named *CBP60A*, *FRS5*, *CRK6*, *PNC1*, *HSP70-16*, *WRKY70*, *TPI*, Thioredoxin H-type 2,and Cytochrome b-c1 complex subunit 7, were found in blue module and in green module of sufficient irrigation (Fig. 5S). The specific genes with high connectivity were RBP45, GLO1, SCPL50, and PIL13 in blue module in severe drought (Fig. 5S–B).

### The relationship between hub genes and relevant genes regulated *g*_m_

Unfortunately, none of the genes regulated *g*_m_ such as aquaporins and cell wall metabolic genes have been discovered in all modules mentioned above. Therefore, protein-protein interaction (PPI) analysis was adopted to explore intrinsic interaction relationships between hub genes in modules in severe drought and the related genes directly regulated *g*_m_ such as aquaporin genes, cell wall regulation genes, and carbonic anhydrase genes. The detailed gene information was presented in Table 2S.The results shown that only genes in midnightblue module had a strong relationship with the genes directly regulated *g*_m_ (Table 3S). Genes of *RBP45*, *GLO1*, *SCPL50*, and *PIL13* in blue module had a poor relationship with the genes directly regulated *g*_m_ (Table 3S). There was little correlation for the other genes in blue module and all genes in the turquoise module with *g*_m_-related genes (Table 3S).

Based on the above results, genes of high connectivity in midnightblue module with *g*_m_-related genes were analyzed using PPI method. Finally, 8 genes (*CSP41B*, *RBCS*, *PGLP1A*, *GAPA*, *DAAT*, *LHCA5*, *CAS*, *GSTU6*) had high correlation with some key genes directly regulated *g*_m_ (Fig. 8). Genes of *DTC*,*PPH*, and *EO* had also high correlation with *g*_m-_related genes, but the genes were not located in the midnightblue module (Table 3S). Moreover, *RhGT1*, *1-sep*, and *SSL4* had low connectivity in midnightblue module, even if those gene had high correlation with *g*_m-_related genes (Table 3S).

**Fig. 8.**
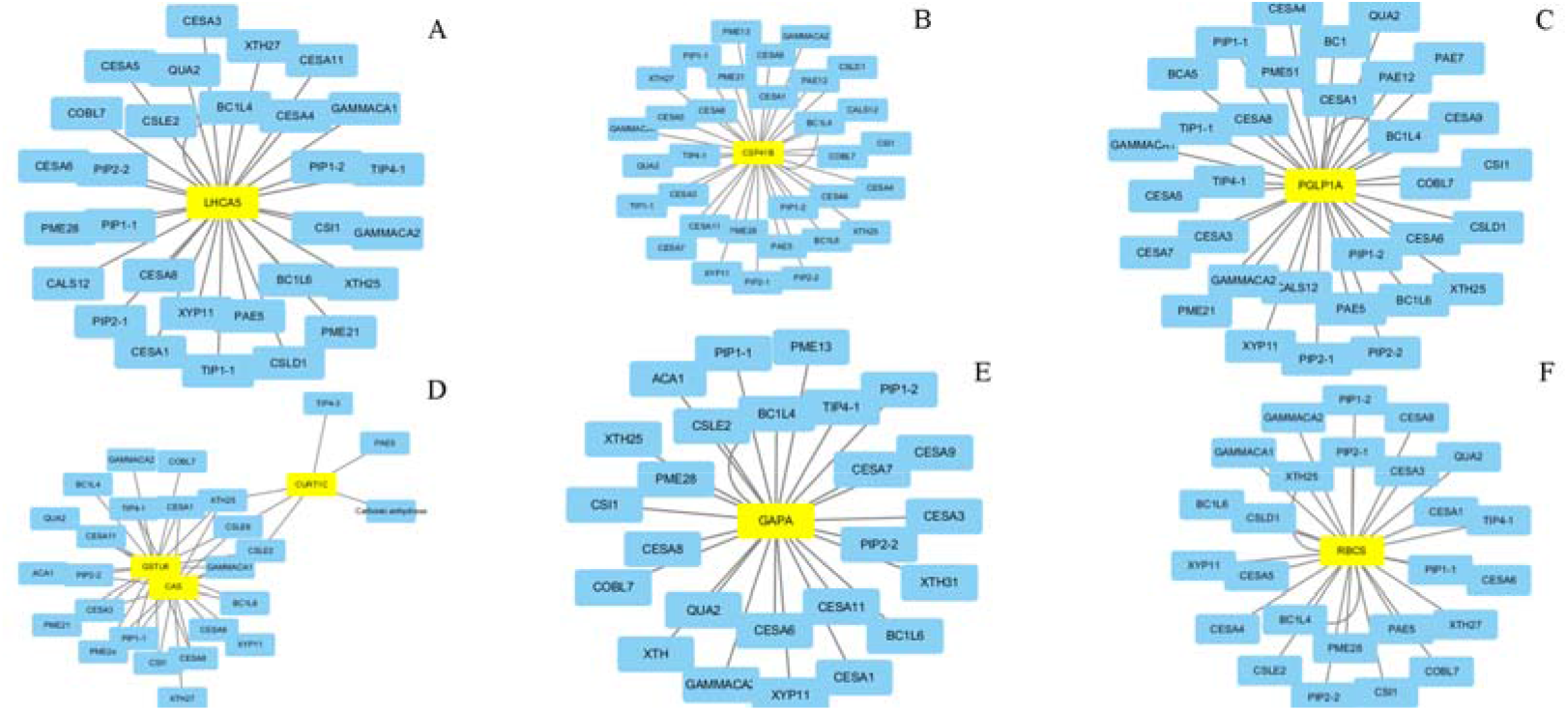
Correlation network diagram between hub genes in the midnightblue module (black letter with yellow background) and the genes directly regulated *g*_m_ (black letter with blue background).

### The relative expression levels of candidate genes among treatments

The genes in midnightblue module and blue module with close relations to the *g*_m_-related genes were considered as candidate genes regulated *g*_m_. 4 genes were from blue module (Fig. 9A–D). And 14 genes came from midnightblue module, but only 11 genes were detected in leaf by RT-PCR method across both cultivars, both observed periods, and three water treatments (Fig. 9E–O). In general, the genes of *RBP45*, *PIL13*, *CSP41B*, *PGLP1A*, *LHCA5*, *GAPA*, *GSTU6*, *CAS*, *PPH*, *SSL4*, and *1-sep* were significantly down-regulated in severe drought compared with sufficient irrigation in both cultivar (Fig. 9; *P*<0.05). However, expression levels in cultivar HY73 had a smaller reduction than that in cultivar HLY898 for the genes of *CSP41B*, *PGLP1A*, *LHCA5*, and *GSTU6* in severe drought when compared with sufficient irrigation (Fig. 9E, F, H, J). Meanwhile, *CSP41B*, *PGLP1A*, *LHCA5*, and *GSTU6* genes had a greater expression levels in cultivar HY73 than that in cultivar HLY898 in both observed periods of three water treatments (Fig. 9E, F, H, J). Especially for the −10 KPa treatment, *CSP41B* and *LHCA5* genes were up-regulation 1.4~2.0 folds in cultivar HY73 in severe drought when compared with cultivar HLY898 in sufficient irrigation (Fig. 9E and H). Only *RBCS* gene was significant up-regulation in severe drought for both cultivars and three water treatments (Fig. 9G; *P*<0.05). The relative expression levels of the *SCPL50* gene were increased by tens of times in severe drought compared with sufficient irrigation for the cultivar HY73 (Fig. 9C; *P*<0.05). But there were no significant difference among three water treatments for the *RBCS* and *SCPL50* genes (Fig. 9C, G; *P>*0.05).

**Fig. 9.**
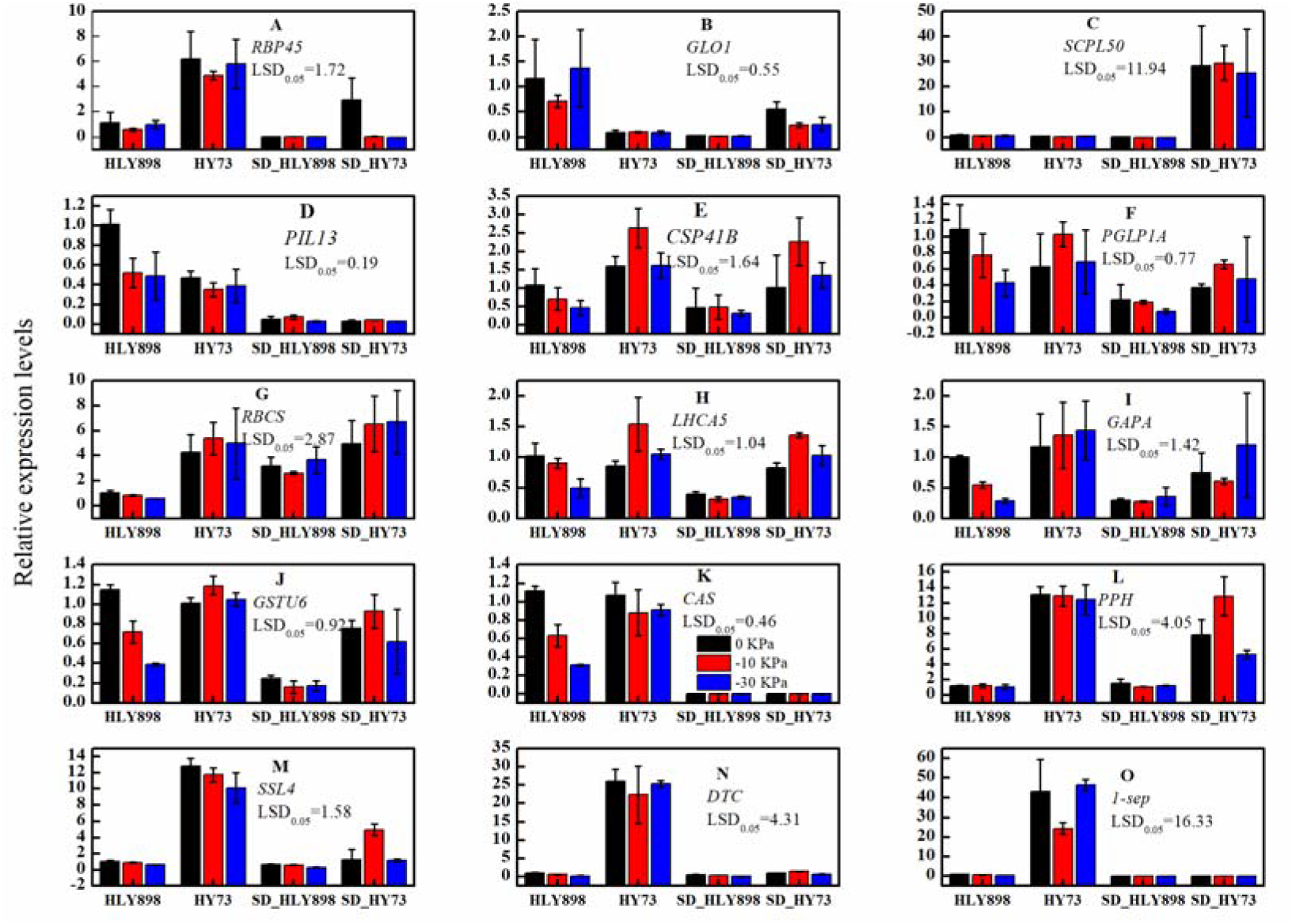
The relative expression levels of genes with a close connection with *g*_m_-related genes in blue module (A-D) and in midnightblue module (E-O) in two observed periods of three water treatments of both cultivars. HLY898 and HY73 indicate the cultivars in sufficient irrigation with three water treatment (0 KPa, −10 KPa, and −30 KPa). SD_HLY898 and SD_HY73 indicate the cultivars in severe drought with three water treatment (0 KPa, −10 KPa, and −30 KPa). The values of LSD _(0.05)_ in each figure are used for multiple comparison of significant differences between both treatments across both cultivars, both observed periods, and three water treatments.

In addition, the relative expression levels of *CSP41B*, *PGLP1A*, *LHCA5*, and *GSTU6* genes had significant linear relationships with *g*_m_ parameter (Fig. 10E, F, H, J). The determinate coefficient (R^2^) were reached to 0.87 for the *PGLP1A* and *LHCA5* genes for the linear fitted equations (*P*<0.00001). There were no any linear relationships between for the rest genes and *g*_m_(Fig. 10).

**Fig. 10.**
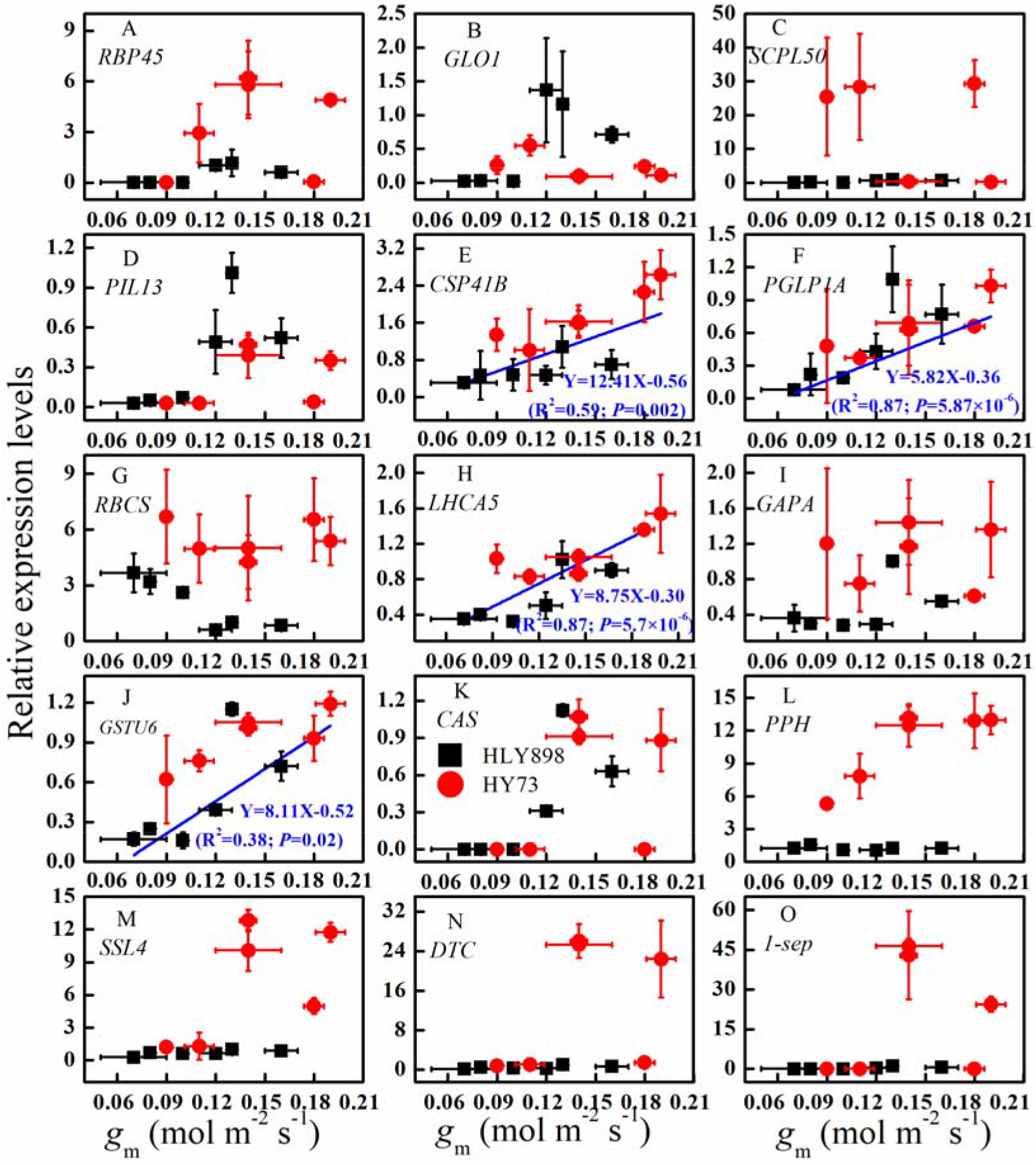
the fitted equations between relative expression levels of candidate genes and *g*_m_ parameter (n=12). For each gene, the data from different water treatments and both cultivars were merged for both cultivar. Black and solid box represent cultivar HLY898. Red and solid circle is cultivar HY73. The bars are standard deviation of *g*_m_ (X-axial) and each gene (Y-axial) from three biological repetitions.

## Discussion

### Transcriptional level traits responded to drought of WDR and it’s relationship with *g*_m_

In the traidtional irrigation, drought-tolerant variety have higher diferentially expressed genes (DEGs) than drought-sensitive variety at the transcriptional level when rice plants are suffered from drought (Ereful *et al*., 2020). Our results were agreement with the view that the number of DEGs was lower for the tolerant cultivar HY73(6149) than for the sensitive cultivar HLY898(6861) when the traidtional irrigation (W1 treatment) was treated with severe drought (Fig. 2).Sobreiro *et al*. (2021) suggest that inert responsive to drought is likely due to morphological and anatomical adaptations to drought. It is meant that cultivar HY73 could have more excellent leaf structural traits than cultivar HLY898 inferred from the results of fewer DEGs in cultivar HY73. This conclusion can be supported by our previous research results. Cultivar HY73 with high photosynthetic production potential is mainly related to well anatomical characteristics to positively regulate *g*_m_ in severe drought when compared with cultivar HLY898 (He *et al*., 2021). In addition, cultivar HLY898 had a higher DEGs than cultivar HY73 in the W1 treatment, but there was a slight difference for the down-regulated differential genes between both cultivars. The differences mainly came from up-regulated differential genes, which cultivar HLY898 had more 575 differential genes than cultivar HY73 (Fig. 2). The results indicated that up-regulated differential genes could play an important role in negative regulation morphological and structural characteristics of leaves when rice plants suffered from severe drought.

Irrigation regimes and fertilizer management are another factors determined the number of DEGs except for cultivars when responded to different environments (Yang *et al*., 2017; Mithra *et al*., 2021). The W2 treatment is considered as an optimized irrigation regime in improving adaptability to severe drought stress of rice plants because W2 treatment have higher photosynthetic physiological performance than the W1 and W3 treatments in severe drought across both cultivars (Fig. 1; He *et al*., 2021). The number of down-regulated differential genes were obvious increase in the W2 treatment compared with the W1 treatment in severe drought (Fig. 2). The results shown that increased larger down-regulated differential genes in the W2 treatment could be beneficial to motivating more metabolic pathways to defend against drought. Then, a higher photosynthetic production ability was obtained in the W2 treatment than in the W1 treatment in the drought (Fig. 2). However, the down-regulated differential genes were then reduced in the W3 treatment for both cultivars when compared with the W2 treatment (Fig. 2). The potential reason would be that long-term drought in the W3 treatment would damage leaf structure and physiological activities regulated photosynthesis process (Ouyang *et al*., 2017; Xiong *et al*., 2017). The metabolic capacities could be further constrained when the W3 treatment was imposed to the environment of severe drought. Finally, the number of down-regulated differential genes were decrease in the W3 treatment (Fig. 2). Meanwhile, the results indicate that down-regulated differential genes could have a positive effects in regulating metabolic processes of cell to maintain high drought resistance for rice plants. Great *g*_m_ of WDR is probably relevant to smaller number of up-regulated differential genes to contribute formation stable leaf structure characteristics and larger number of down-regulated differential genes to maintain high metabolic activities in severe drought compared with drought-sensitive cultivar.

### Physiological mechanisms regulated *g*_m_ in WDR

*g*_s_ and *g*_m_ are the main restricted factors to leaf photosynthesis of rice in adequate irrigation and drought conditions (Ikawa *et al*., 2019; Tanaka *et al*., 2019). In sufficient irrigation, 2370 genes had significant relationships with *g*_s_ and *P*_n_ (*P*<0.05). Only 81 genes were related to *g*_m_, but there were no correlational relationship between the genes and *P*n (Fig. 3A). The results showed that the effects of 2370 genes on *P*_n_ might be mainly affected by stomatal pathway. Previous research results, including our study, also support this view that high *P*_n_ needs to match great *g*_s_ for rice plants in sufficient irrigation (Fig. 1 A and B; Ikawa *et al*., 2019). In severe drought, decreasing amplitude of *g*_m_ was lower than that of gs compared with sufficient irrigation for three water treatments of both cultivars. The trends of *P*_n_ was basically consistent with *g*_m_ among water treatments for both cultivars (Fig. 1). The results could indicate that g_s_ is more sensitive to severe drought than *g*_m_ and *g*_m_ is main factor to regulate *P*_n_ in severe drought of rice plants. The founds are basically consistent with previous studies that maintaining high *g*_m_ is key to obtain high *P*_n_ in drought (Ouyang *et al*., 2017; He *et al*., 2021; Li *et al*., 2021).

Interestingly, clustered 3071 genes assigned to midnightblue, blue, and turquosie modules had significant relationship with *g*_m_ in severe drought (*P*<0.05; Fig. 3B). No gene was found a significant correlation with *g*_s_ in severe drought (*P*>0.05; Fig. 3B). These results imply that at least 3071 genes could be involved in regulating *g*_m_ by direct and indirect ways to impede its significantly decrease in severe drought. In the three modules, only midnightblue module with 734 genes played direct functions to manage *g*_m_ (Fig. 6; Table 2). It is widely accepted that aquaporin, carbonic anhydrase, cell wall traits, and chloroplast structure directly control the CO_2_ diffusion capacities from intercellular space to carboxylation site of chloroplast, i.e. *g*_m_ (Flexas *et al*., 2012). But unfortunately, no gene were involved to translate the processes in the 734 genes (Fig. 6; Fig. 7;Table 2). Main function of the 734 genes were annotate in PSII and PSI systems (Table 2). As we known, PSII and PSI system are located on the thylakoid membrane (Wang *et al*., 2019b), which the main contribution to photosynthetic carbon fixation is formation energy such as NADPH and ATP by quantum absorption and electron transport in PSII and PSI system (Kirchhoff, 2018). Energy distribution traits between ATP and NADPH is pivotal to link light reaction, CO_2_ assimilation and other metabolism pathways (Cardona *et al*., 2018; Wang *et al*., 2019a). Therefore, high photosynthetic production formation could be related to optimized energy distribution, especially during NADPH and ATP yielding process in PSII and PSI system in thylakoid membrane, except for maintaining high *g*_m_ in severe drought in WDR. However, the interaction mechanisms of *g*_m_ coupling energy distribution in PSII and PSI system is unclear in drought of rice plants.

Some theories can support this hypothesis that *g*_m_ coupling energy distribution in PSII and PSI system could be key factors to synergistically manage photosynthetic production potential. When CO_2_ concentration at carboxylase site is restricted by low g_s_ or *g*_m_ in adverse environmental conditions, the amount of NADP^+^, which was regenerated in tricarboxylic acid cycle processes of photosynthesis, would be limited because of inadequate CO_2_ supply capacities. And then adjusting energy redistribution in PSII and PSI systems to avoid photoinhibition and photo-damage of the both photosystems (Zivcak *et al*., 2013; Li *et al*., 2021). In addition, low *g*_m_ in stress restrain carboxylation rate and regeneration of ribulose diphosphate, electron transport in PSII is impaired in the circumstances (Killi and Haworth, 2017). Therefore, the high photosynthetic production potential of WDR in severe drought would be fully understood by revealing the interaction mechanisms between *g*_m_ and energy distribution in PSII and PSI system. Mesophyll cell structures including anatomical structure, ultrastructures of chloroplast and thylakoid membrane, and intracellular physiological traits such as aquaporins, carbonic anhydrase, and thylakoid membrane proteins and protein complexes have different functions and contribution pathways to regulate *g*_m_ and energy distribution during responding to stress environment in rice plants (Flexas *et al*., 2012; Xiong *et al*., 2017; Wang *et al*., 2018b; Pashayeva *et al*., 2021; Wu *et al*., 2021). However, it is still indistinct how the cell structures interacted with physiological functions regulate *g*_m_ and energy distribution to maintaining high photosynthetic production ability in water stress in rice plants.

### Molecular mechanisms regulated *g*_m_ in WDR

In our study, genes that directly regulate *g*_m_, for instance, PIPs and TIPs, carbonic anhydrase synthesis gene, and some of genes associated with cell wall metabolisms, had not been discovered in drought condition from hub gene pools of all modules (Fig. 7). Interestingly, four candidate genes named LHCA5, CSP41B, GSTU6 and PGLP1A had a similar tendency among different treatments of both cultivars compared with *g*_m_ (Fig. 9). Treatments with great *g*_m_ generally presented a high relative expression level for the genes. There were significant linear relation between gm and the genes merged the data of cultivars, water treatments, and both observed periods (Fig. 10E, F, H, and G). Moreover, LHCA5 and CSP41B genes were significantly up-regulated after applying severe drought (Fig. 10E and H). Therefore, LHCA5 and CSP41B could play a positive functions regulating *g*_m_.

LHCA5 gene is usually expressed in the nucleus predicted by subcellular localization methods and it highly conserved across different crops (Kato *et al*., 2018). Existed studies have confirmed that LHCA5 is an important component of the chloroplast NADH dehydrogenase supercomplex (NDH), which plays an important role in regulating cyclic electron flow (Otani *et al*., 2018). In severe drought, the expression level was up-regulated for the LHCA5 gene for WDR when compared with cultivar HLY898, especially in the W2 treatment (Fig. 9H). The results shown that cyclic electron flow rate could be enhanced for WDR in severe drought compared with cultivar HLY898. Cyclic electron transport can produce energy ATP, which the process is considered as a effective ways to avoid excessive reduction of receptor side in PSI system (Christine and Shigeru, 2011). In addition, the ATP generated within cyclic electron flow process is mainly used for consuming in photo-respiration metabolic for crops (Yi *et al*., 2018; Wang *et al*., 2019a). Therefore, photo-respiration could be higher in severe drought in WDR than in cultivar HLY898 because WDR had greater cyclic electron flow rate. The results also indicated that high *g*_m_ in WDR could be benefits from good light adaptation mechanisms in drought by up-regulating gene express of LHCA5 gene compared with HLY898.

Moreover, LHCA5 was closely related to the genes regulated *g*_m_ such as aquaporin genes, carbonic anhydrase genes, cellulose synthase genes, and syloglucan endoglycosidase/hydrolase genes. Meanwhile, LHCA5 gene is bound up with light-absorbing pigment molecule LHCA1-4 (Ulrika *et al*., 2004). Therefore, LHCHA5 may play an important role in regulating energy distribution in photosystems by manipulating the absorption capacitied of light quanta and electron transport pathway except for affecting *g*_m_. In other words, LHCA5 gene may be a key functional gene in co-regulation energy distribution in photosystems and *g*_m_. However, the molecular mechanisms of OsLHCHA5 on energy distribution in photosystems and *g*_m_ has not been reported yet.

CSP41B gene had localized in chloroplasts in recent year (Mei *et al*., 2017). The function of the gene is related to leaf color (Thomas *et al*., 2009; Mei *et al*., 2017). CSP41B is a required gene for normal leaf color (Mei *et al*., 2017). High express of CSP41B have a positive function to maintaining heavy green leaves of crops (Thomas *et al*., 2009; Mei *et al*., 2017). In severe drought, WDR had higher express level than cultivar HLY898 for CSP41B gene. It is meant that leaf color was more light-green color in cultivar HLY898 than that in WDR in severe drought. The phenomenon can be supported by our observation for SPAD value that WDR had higher SPAD value than cultivar HLY898 in severe drought, especially for the W2 treatment (Data not presented in here). The results shown that aging phenomenon aggravate for cultivar HLY898 compared with WDR in severe drought. Then, photosynthesis would be down-regulated because of inadequate energy capture and transfer efficiency in low pigment molecular content in cultivar HLY898. Furthermore, CSP41B gene has also positive function in regulating chloroplast morphology (Mei *et al*., 2017). The function might be useful to explain the results that the CSP41B gene were closely related to the gene that regulates mesophyll conductance. CSP41B could play an important role affecting *g*_m_ and energy absorption, transport, and distribution, but there are still not clear how the CSP41B gene synergistically regulate them.

## Conclusion

i. For WDR, up-regulated differential expression genes (DEGs) are lower than drought-sensitive variety in severe drought. But the number of down-regulated DEGs are significantly higher in WDR than in drought sensitive variety. DEGs had an obvious increased trend from pre-slight drought irrigation regime to pre-mild drought irrigation regime before severe drought in WDR. However, DEGs was sharply decreased in pre-mild drought irrigation regime before severe drought in drought-sensitive variety. The results suggest that there are different regulation mechanisms to *g*_m_ at transcriptional level for WDR and drought sensitive variety. In general, high drought-resistance ability in WDR could be related to greater endurance by activating more differential genes when respond to complex environment compared with drought sensitive variety.
ii. Function of thylakoid membrane is closely related to *g*_m_. The formation of high photosynthetic production potential may be the result of the synergistic effects of gm and energy distribution in thylakoid membrane for WDR in severe drought.
iii. *OsLHCA5* and *OsCSP41B* genes locate in thylakoid membrane and regulate energy absorption, transport and distribution functions. The two genes have positive relationship with the genes that directly regulated gm. Therefore, *OsLHCA5* and *OsCSP41B* genes could be candidate genes for the synergistic regulation of *g*_m_ and energy allocation in thylakoid membrane to obtain high photosynthetic production potential in drought.

## Supplementary Information

Table 1S. Primer sequence of candidate genes

Table 2S. Genes directly regulated *g*_m_ (The table have been uploaded to the submission system)

Table 3S. Correlation between genes in each module and the genes directly regulated *g*_m_ (The big table table have been uploaded to the submission system)

## Author contributions

Conceived and designed the experiments: Haibing He, Liquan Wu. Performed the experiments: Lele Wang, Quan Wang. Analyzed the data: Haibing He, Xuelan Zhang, Ru Yang. Contributed reagents/materials/analysis tools: Li Zhan, Cuicui You, Jian Ke. Wrote the manuscript: Haibing He. Corrected the manuscript: Liquan Wu.

## Availability of Data and Materials

All data supporting the conclusions of this article are provided within the article.

## Declarations

Ethics approval and consent to participate: Not applicable.

Consent for publication: Not applicable.

---

**Fig .1S.**
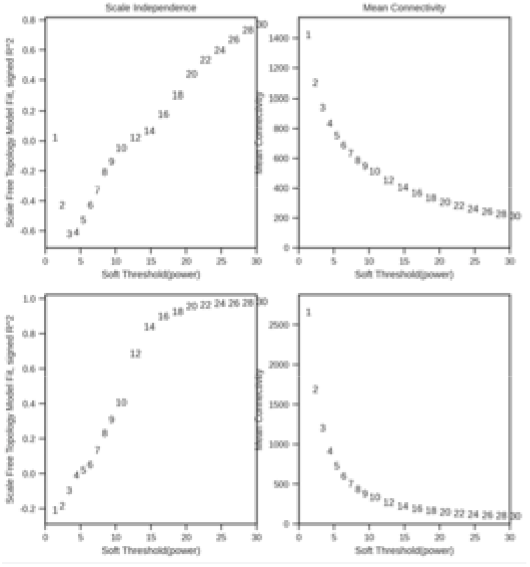
R^2^ of soft threshold (A and C) and mean connectivity (Band D) of gene co-expression network in sufficient irrigation (A and B) and in severe drought (C and D). When soft threshold were 30 and 14 in sufficient irrigation and severe drought, respectively. R^2^>0.8 at both observed stages.

**Fig. 2S.**
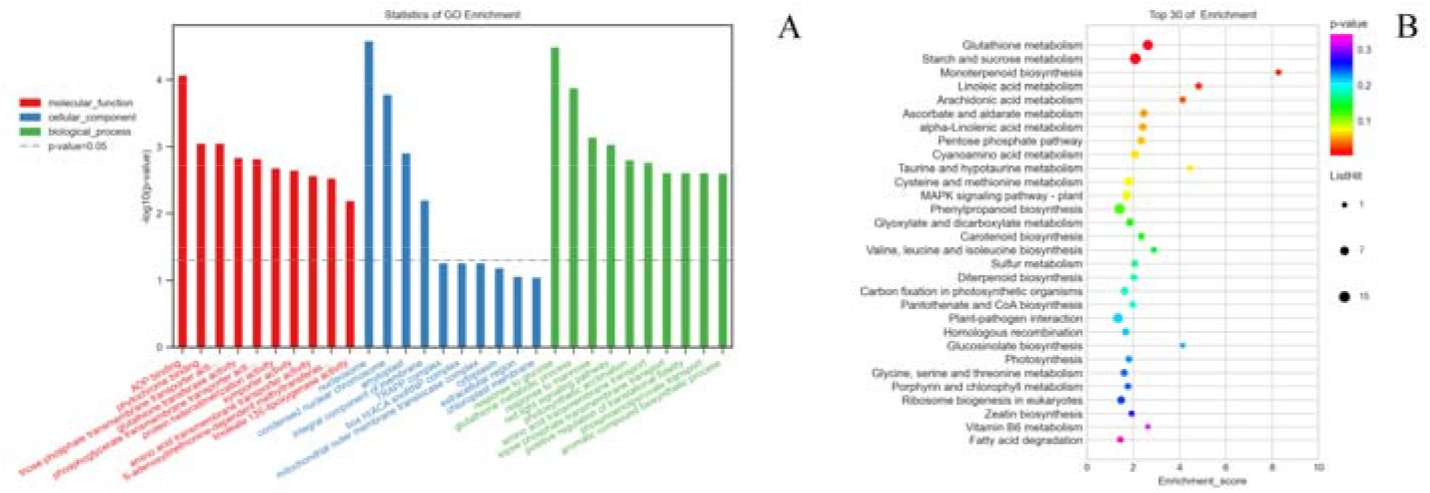
The first 10 GO (A) and 30 KEGG (B) items in the brown module based on WGCNA analysis 350 in sufficient

**Fig. 3S.**
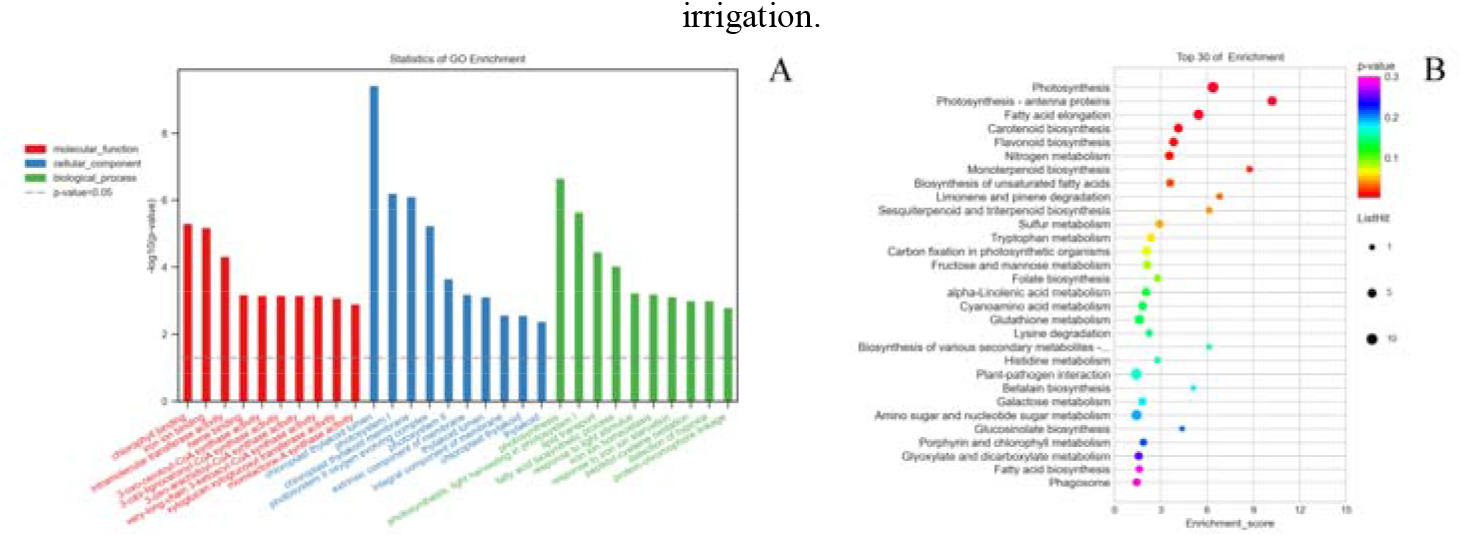
The first 10 GO (A) and 30 KEGG (B) items in the green module based on WGCNA analysis in sufficient

**Fig. 4S.**
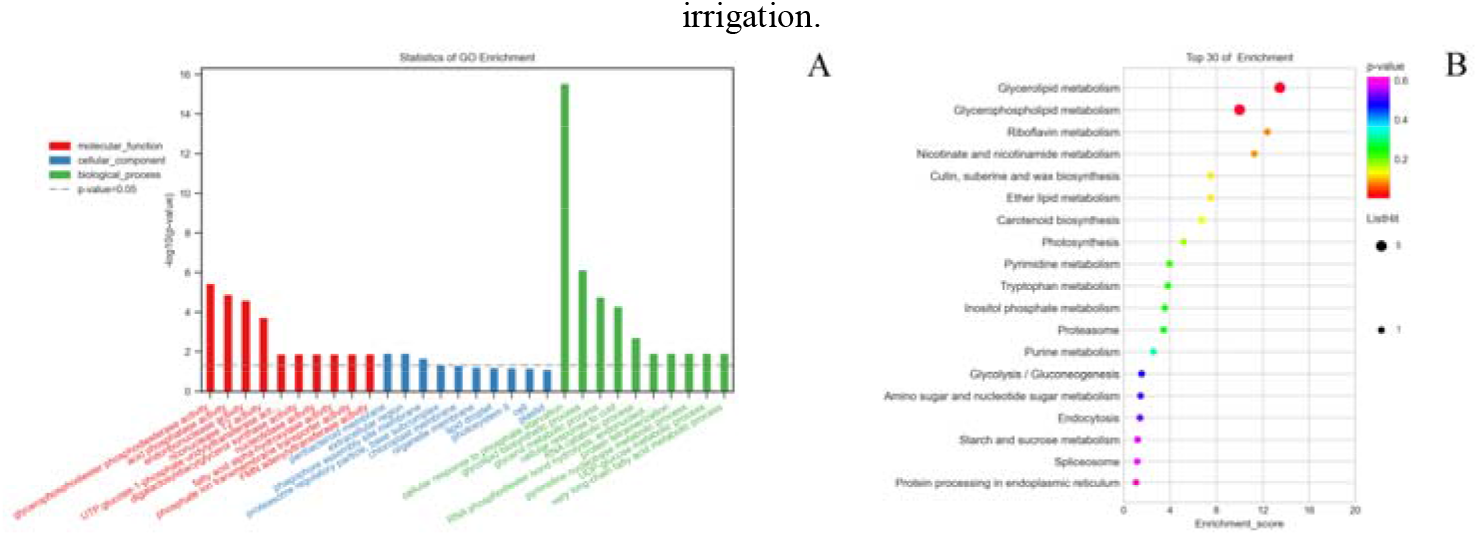
The first 10 GO (A) and 30 KEGG (B) items in the red module based on WGCNA analysis in sufficient irrigation.

**Fig. 5S.**
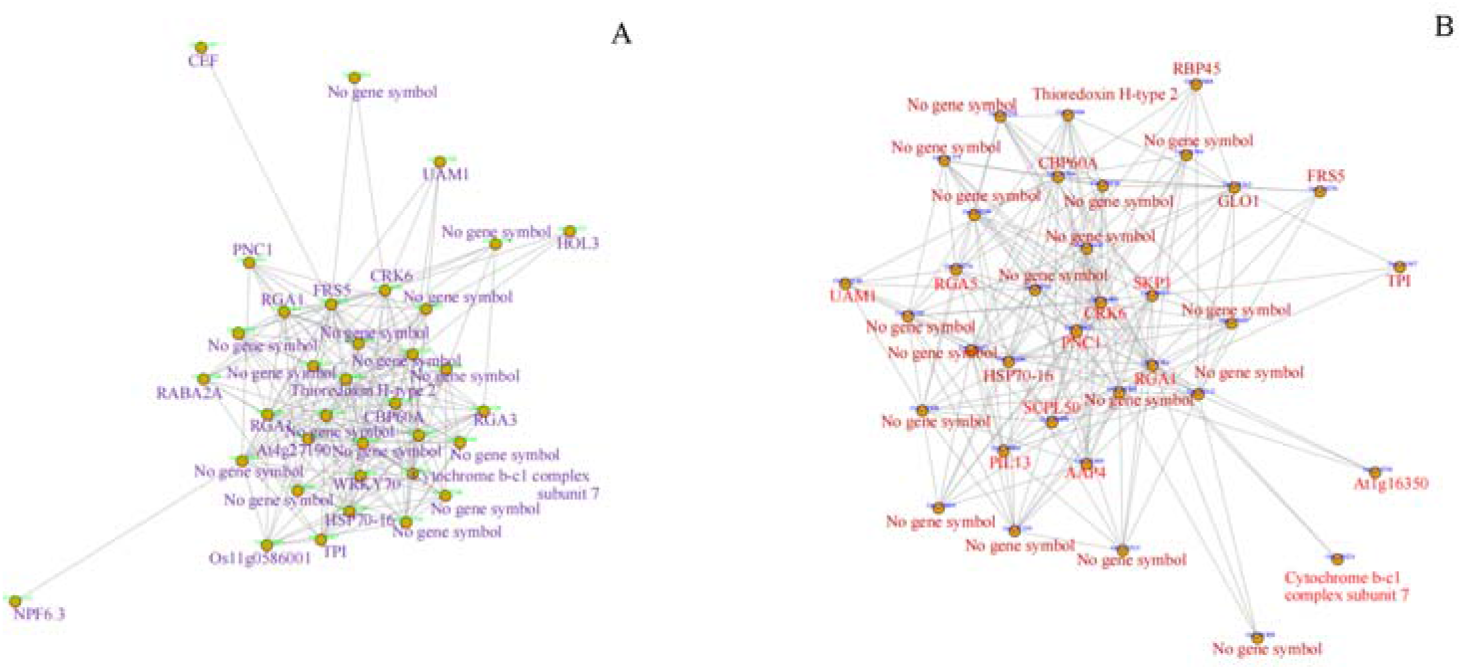
Co-expression networks of the most highly connected hub genes in the green module in sufficient irrigation (A) and in the blue module in severe drought (B).

**Fig. 6S.**
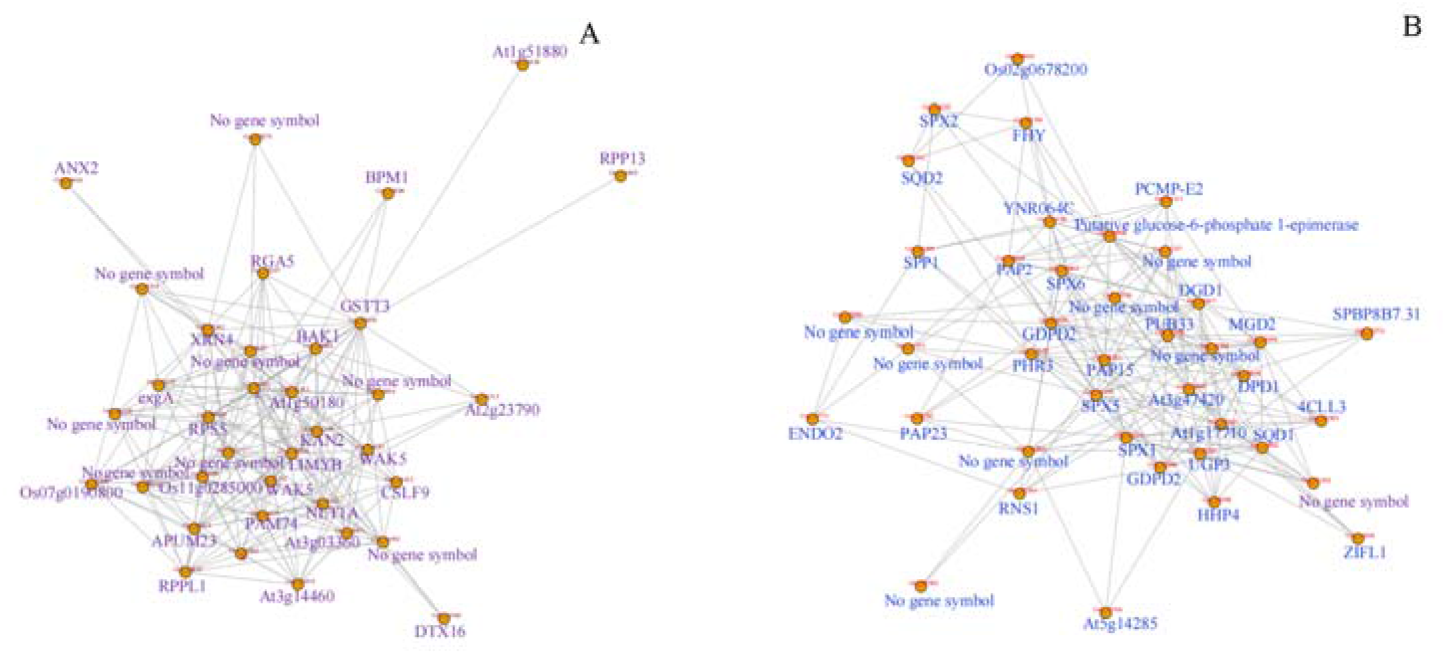
Co-expression networks of the most highly connected hub genes in the brown (A) and red (B) modules in severe drought.

